# Dynamic Brain Connectivity Patterns in Conscious and Unconscious Brain

**DOI:** 10.1101/075788

**Authors:** Yuncong Ma, Christina Hamilton, Nanyin Zhang

**Affiliations:** Department of Biomedical Engineering, Pennsylvania State University, University Park, 16802; The Huck Institutes of Life Sciences, Pennsylvania State University, University Park, PA, 16802

**Keywords:** anesthetic-induced unconsciousness, dynamic functional connectivity, resting-state networks, rat

## Abstract

Brain functional connectivity undergoes dynamic changes from the awake to unconscious states. However, how the dynamics of functional connectivity patterns are linked to consciousness at the behavioral level remains elusive. Here we acquired resting-state functional magnetic resonance imaging (rsfMRI) data during wakefulness and graded levels of consciousness in rats. Data were analyzed using a dynamic approach combining the sliding-window method and k-means clustering. Our results demonstrate that whole-brain networks contain several quasi-stable patterns that dynamically recurred from the awake state into anesthetized states. Remarkably, two brain connectivity states with distinct spatial similarity to the structure of anatomical connectivity were strongly biased toward high and low consciousness levels, respectively. These results provide compelling neuroimaging evidence linking the dynamics of whole-brain functional connectivity patterns and states of consciousness at the behavioral level.

**Conflict of interest:** none.

## Introduction

Brain function undergoes dramatic changes from conscious to anesthetic-induced unconscious (AIU) states. Changes in brain function during AIU are intrinsically dynamic, considering that consciousness is known to be not an all-or-none, but a graded phenomenon (Laureys et al., 2007). However, what these dynamics are, and how dynamical changes in brain function are linked to consciousness at the behavioral level, remain unknown.

In recent years, accumulating evidence suggests that AIU is tightly associated with functional connectivity (FC) changes in brain networks (Lee et al., 2013; Liang et al., 2013; Mashour and Alkire, 2013; White and Alkire, 2003). These functional brain networks have been uncovered by neuroimaging techniques, especially resting-state functional magnetic resonance imaging (rsfMRI) (Boveroux et al., 2010), which measures resting-state functional connectivity (RSFC) based on temporal synchronizations of blood-oxygenation-level dependent (BOLD) signals from different brain regions (Biswal et al., 1995; Fox and Raichle, 2007). Although rsfMRI has been extensively used to study brain function during AIU, the conventional correlation analysis that rsfMRI uses implicitly assumes that RSFC is stationary during data acquisition. Therefore, in essence, such analysis only provides averaged RSFC over the recording period, while the information of how RSFC temporally varies is overlooked. Given this nature, the most majority of previous rsfMRI studies focused on identifying steady-state connectivity changes between two or three anesthetic depths (Liang et al., 2012b; Vincent et al., 2007).

Despite the tremendous progress the steady-state rsfMRI paradigm has made, it may not be optimal to reveal the dynamic changes of brain network connectivity when the consciousness level is modulated. This limitation has highlighted a critical knowledge gap as the dynamics of whole-brain network connectivity at and in transition to different levels of consciousness will help decipher network reconfigurations when consciousness is perturbed. Therefore, this information can provide critical insight into understanding the systems-level neural mechanisms of AIU.

Recent progress in rsfMRI methodology demonstrates that dynamic information of brain connectivity can be reliably detected in humans and animals with novel analysis methods such as the sliding-window and co-activation pattern approaches (Allen et al., 2014; Chang and Glover, 2010; Hutchison et al., 2013a; Hutchison et al., 2013b; Keilholz et al., 2013; Liang et al., 2015a; Liu and Duyn, 2013; Thompson et al., 2013; Thompson et al., 2014). These methods have made it feasible to investigate the brain network dynamics at different states of consciousness (Barttfeld et al., 2015; Liang et al., 2015a). Indeed, with these approaches, it has been shown that the dynamic properties of brain network connectivity may constitute key characteristics of consciousness (Barttfeld et al., 2015; Breshears et al., 2010; Hudetz et al., 2015).

To comprehensively elucidate the dynamics of brain connectivity patterns in different states of consciousness, here we acquired rsfMRI data in rats during wakefulness and five graded levels of consciousness induced by increasing concentrations of isoflurane. The animal’s state of consciousness was assessed using the loss of righting reflex (LORR) test. rsfMRI data were analyzed using the sliding-window method combined with k-means clustering (Allen et al., 2014). We identified two specific brain network patterns that were strongly associated with high and low consciousness states, respectively. Interestingly, we also found a brain connectivity state that was dominant at the critical transitional point between the conscious and unconscious states. Based on these findings, we further analyzed the temporal sequences of separate connectivity patterns between conscious and unconscious states.

## Methods and Materials

### Animal Preparation

All procedures were conducted in accordance with approved protocols from the Institutional Animal Care and Use Committee of the Pennsylvania State University. 25 adult male Long-Evans rats (300-500g) were housed and maintained on a 12hr light: 12hr dark schedule, and provided access to food and water *ad libitum* throughout the duration of the study. To minimize stress and motion during imaging at the awake state, animals were acclimated to the scanning environment for 7 days (described in (Liang et al., 2011, 2012a, 2014; Liang et al., 2013; Liang et al., 2015b; Zhang et al., 2010)). To do this, rats were restrained and placed in a mock MRI scanner where prerecorded MRI sounds were played. The exposure time in the mock scanner started from 15 minutes on day 1, and was incrementally increased by 15 minutes each day up to 60 minutes (days 4, 5, 6, 7). This setup mimicked the scanning conditions inside the magnet.

### Behavioral Assessment

The animal’s consciousness state was assessed using the behavioral test of LORR. A strong correlation has been shown between the concentration of various anesthetic agents necessary for rodents to lose voluntary movement and loss of consciousness in humans (Franks, 2008). In the LORR test, each rat began in the awake state in the restrainer and was exposed to increasing doses of isoflurane through a nosecone. The LORR for a single dose was measured exactly as it was delivered to the animal inside the scanner (Fig. 1A). Each rsfMRI scan was 10-minute long with a 5-minute inter-scan transition time. Thus, for instance, LORR at 1.5% isoflurane was measured by sequentially delivering 0.5% and 1% isoflurane to the animal for 15 minutes, respectively, followed by 1.5% isoflurane for 5 minutes. With the nosecone in place, the rat was then removed from the restrainer and placed in a supine position while the time it took to correct its position was measured. The animal was deemed completely unconscious if it did not correct its position within 60 seconds. This procedure mimicked anesthesia administration during the rsfMRI scanning process, and thus allowed us to verify the animal’s consciousness state during each rsfMRI scan.

**Figure 1.**
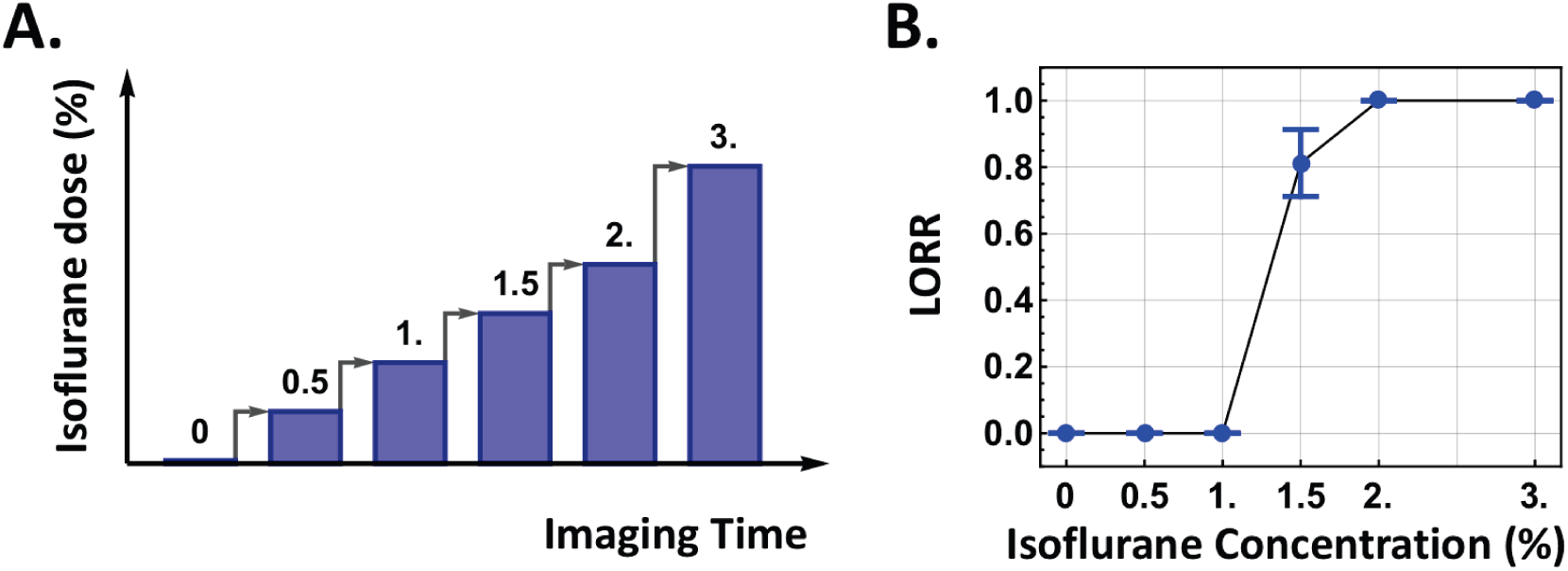
Stationary RSFC strength was reduced during AIU, albeit the spatial pattern of whole-brain network remained similar. rsfMRI paradigm and behavioral test. (A) The imaging paradigm consisted of six sequential rsfMRI scans with an increasing dose of isoflurane. Each 10-min scan was preceded by a 5-min inter-scan interval. (B) The fraction of the number of animals that lost righting reflex at each dose. Bars: S.E.M.

### MRI Experiment

Rats were briefly anesthetized with isoflurane while they were placed in a head restrainer with a built-in birdcage coil. Isoflurane was discontinued after the animal was set up, but the nosecone remained in front of the animal’s nose for the duration of the whole MRI experiment. Imaging began 30 minutes after rats were placed in the scanner while animals were fully awake. Image acquisition was performed on a 7T scanner interfaced with a Bruker console (Billerica, MA). First, a T1-weighted structural image was acquired with the following parameters: repetition time (TR) = 2125 ms, echo time (TE) = 50 ms, matrix size = 256 × 256, field of view (FOV) = 3.2 cm × 3.2 cm, slice number=20, slice thickness = 1 mm. Subsequently, six rsfMRI scans were acquired: one as animals were fully awake and five at increasing isoflurane concentrations delivered at 0.5%, 1.0%, 1.5%, 2.0% and 3.0%, respectively, with a 5-minute transition time between scans (Fig. 1A). Isoflurane was delivered via the nose cone. rsfMRI data were acquired using a single-shot gradient-echo echo planar imaging pulse sequence with TR = 1000 ms, TE = 13.8 ms, flip angle = 80°, matrix size = 64 × 64, FOV = 3.2 × 3.2 cm^2^, slice number = 20, 1mm thick slices (in-plane resolution = 0.5 ×0.5 mm^2^). 600 rsfMRI volumes (10 min) were acquired for each scan (Figure 1).

All imaging and behavioral data were also used in another study (Hamilton et al., 2016) and reanalyzed for the purpose of this paper.

### Data Analysis

#### Image Preprocessing

The first 10 volumes of each rsfMRI scan were removed to allow magnetization to reach a steady state. rsfMRI data were then manually co-registered to a standard rat atlas using Medical Image Visualization and Analysis software (MIVA, http://ccni.wpi.edu), motion corrected using SPM 12 (University College London, UK), and then spatially (Gaussian kernel, sigma = 0.5mm) as well as temporally smoothed (0.01–0.1Hz). In addition, signals from white matter and ventricles, six motion parameters estimated from motion correction (translation and rotation), as well as their derivatives were regressed out (Power et al., 2015).

#### Brain parcellation

The whole brain was parcellated in to 134 unilateral regions of interest (ROIs) based on the anatomic definition of brain regions in the Swanson Atlas (Swanson, 2004). 110 of these ROIs were further grouped into 9 brain systems including sensory-motor, polymodal association, retrohippocampal regions, hippocampus, amygdala complex, striatum, pallidum, thalamus and hypothalamus (Liang et al., 2012b) (Fig. 2A). The remaining 24 ROIs were noted as ‘others’. The ROI time course was extracted by regionally averaging rsfMRI time series from all voxels within the ROI.

**Figure 2.**
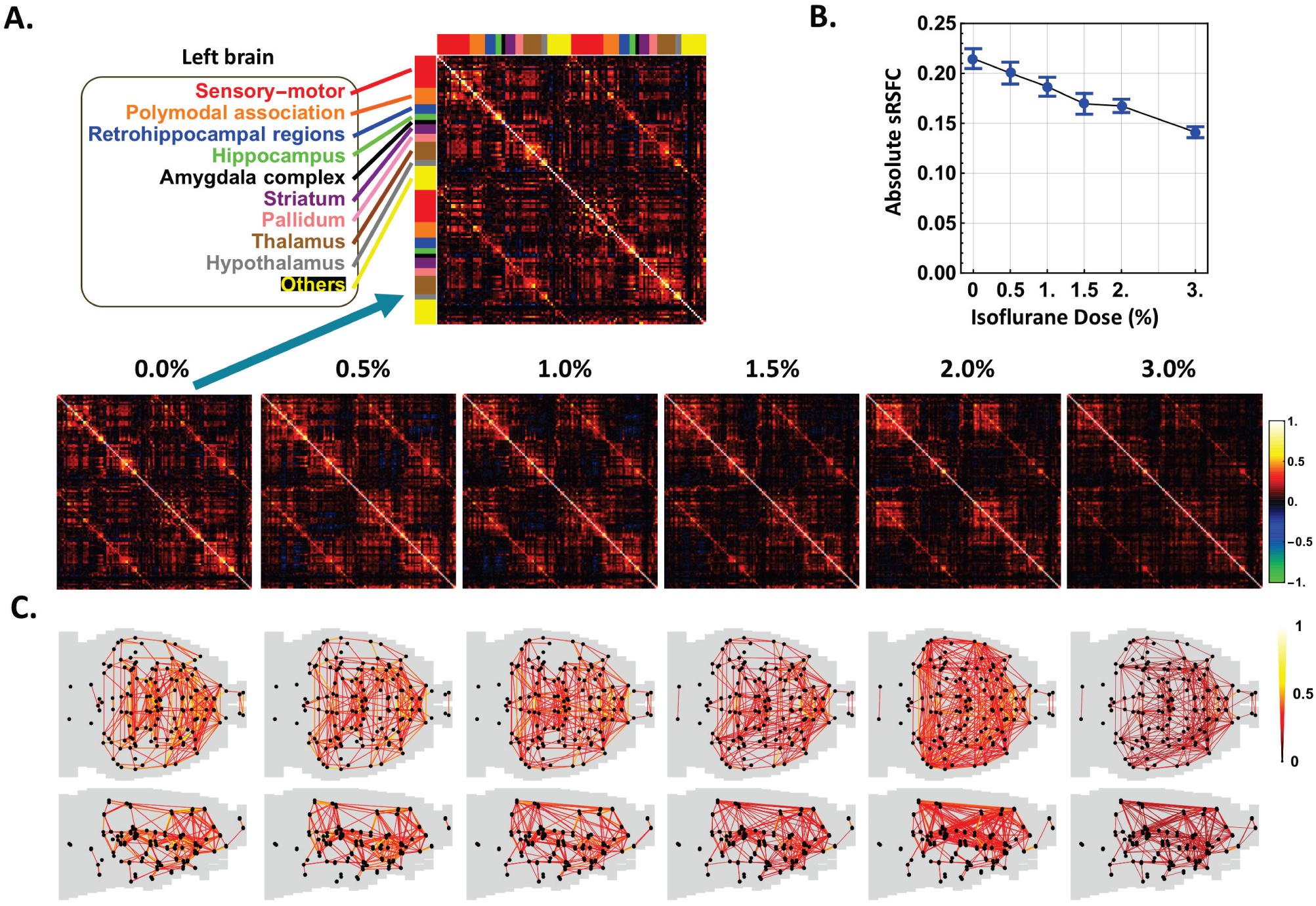
Whole-brain connectivity patterns contained several quasi-stable states that dynamically alternated from the awake state into anesthetized states. sRSFC patterns. (A) sRSFC matrices at the awake state (0.0%) and five anesthetic depths induced by increasing concentrations of isoflurane (0.5%, 1.0%, 1.5%, 2.0% and 3.0%). ROIs were grouped by the hemisphere and brain system (first half: left hemisphere, second half: right hemisphere). Red: sensory-motor system, orange: polymodal-association system, blue: retrohippocampal regions, green: hippocampus, black: amygdala complex, purple: striatum, pink: pallidum, brown: thalamus, gray: hypothalamus, yellow: all other ROIs. (B) Average absolute sRSFC strength during wakefulness and five isoflurane concentrations, bars: S.E.M. (C) Axial and sagittal views of the brain connectivity pattern for all conditions. Dark dots indicate ROIs. For each dose, 446 strongest connections (i.e. 0.05 graph density) were displayed.

#### Estimating anatomical connectivity based on axonal projection density

We used the Allen Mouse Brain Connectivity Atlas to generate the structure of anatomical connectivity between ROIs based on axonal projection density. The Allen Brain Atlas was used due to its’ public availability and the high anatomical and functional similarity between rat and mouse brains (Oh et al., 2014). Considering that the Allen mouse’s anatomical map contains finer brain structures that cannot be differentiated by our fMRI’s spatial resolution, we first combined all small sub-structures in the Allen Brain Atlas that belonged to each ROI defined in the rat brain. Secondly, the average axonal projection density from ROI A to ROI B was assessed by taking the weighted average of the projection densities across all sub-structures in A to all sub-structures in B, with the size of each sub-structure in B as the weight. Mathematically, the average projection density from A to B was defined as:

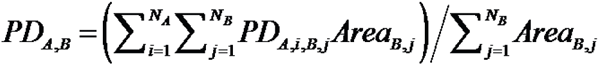

where *PD*_*A,i,B,j*_ is the projection density from the i-th sub-structure in ROI A to the j-th sub-structure in ROI B; *Area*_*B,j*_ is the size of the j-th sub-structure in ROI B; *N*_*A*_ and *N*_*B*_ are the number of sub-structures in ROI A and B, respectively. Finally, the structural connectivity strength between ROI A and B was calculated as the mean of *PD*_*A,B*_ and *PD*_*B,A*_.

#### Deriving brain functional connectivity

We first calculated the brain functional connectivity patterns under the stationarity assumption. The ROI-wise RSFC matrices (referred to as sRSFC matrices hereafter) were obtained by calculating the Pearson correlation coefficients between each pair of ROI time courses at the awake and five anesthetic depths. These functional connectivity matrices were organized by the brain systems defined above (Fig. 2).

Dynamic brain connectivity patterns in different anesthetized states were analyzed using the sliding window method (window length = 60s, step size = 5s, TR = 1 s). A 60-s window size was selected based on the report that window sizes around 30-60 s produce robust RSFC results (Hutchison et al., 2013a; Jones et al., 2012; Shirer et al., 2012). For each rsfMRI scan, 61 windows were extracted for each dose. The RSFC matrix of each window was calculated in the same manner as sRSFC matrices. RSFC matrices for all windows in the awake state and five anesthetic depths from all animals were then pooled together and clustered into five temporarily alternating, but spatially repeatable whole-brain connectivity patterns (referred to as dRSFC hereafter) using k-means clustering (Manhattan distance). These dRSFC patterns were ranked according to their spatial similarity (measured by Manhattan distance) to the structural connectivity map obtained as mentioned above. To facilitate the comparison between dRSFC and structural connectivity, an arctangent function was applied to normalize the dRSFC strength and anatomical projection intensity, respectively. For each dRSFC pattern, the occurrence rate was measured for each consciousness state.

To explore the temporal transitions between dRSFC patterns, the state sequence of each scan, which represents the sequence of rsfMRI windows with the identity of their corresponding dRSFC patterns, was obtained from the clustering results. The number of transitions from one state to another state, which was apart by one window length (i.e. 60 s), was counted. The 60-s gap ensures the temporal independence of the two brain connectivity states in transition counts. The state transition probability from one state to another was then calculated after excluding all self-transitions.

#### Statistics

Isoflurane dose effects were statistically tested using Chi-square test or One-way ANOVA. The relationship between dynamic RSFC patterns and consciousness levels were tested using two-way ANOVA. The comparison between high and low consciousness levels was conducted using two-sample t-tests. The statistical significance was set at p<0.05.

## Results

In the present study, we examined the anesthetic-induced dynamic changes in the whole-brain network connectivity as animals transitioned from the awake state into an unconscious state. rsMRI data during wakefulness and graded levels of consciousness produced by increasing concentrations of isoflurane (0.5%, 1.0%, 1.5%, 2.0% and 3.0%) were collected in rats.

### Assessing the consciousness level at different anesthetized states

To understand the dynamics of network connectivity at different consciousness levels, we determined the animal’s conscious state at each isoflurane dose delivered. The animal’s consciousness level was measured using LORR outside of the scanner in a manner that directly mimicked the way the dose was delivered during rsfMRI scanning (Fig. 1A, also see **Methods**). Our data showed that no rats lost righting reflex when 1% or 0.5% isoflurane was delivered via a nosecone, while 81% of animals were unable to correct from the supine to prone position when 1.5% isoflurane was delivered. All animals lost righting reflex at higher delivered doses (2% and 3%, Fig. 1B). Therefore, loss of consciousness was induced when 1.5% isoflurane was delivered through a nosecone (Chi-square test, χ^2^=22.82, p<2×10^−6^).

We first examined the steady-state brain functional connectivity under the stationarity assumption. The whole brain was parcellated into 134 unilateral ROIs (Swanson, 2004). The ROI-wise RSFC matrices (referred to as sRSFC matrices) were obtained by calculating the Pearson correlation coefficients between pairs of ROI time courses of each rsfMRI scan at the awake state and five anesthetic depths. Consistent with previous studies (Liang et al., 2012b; Liu et al., 2011; Lu et al., 2007; Peltier et al., 2005), sRSFC decreased at increasing anesthetic dose (Figs. 2A, B; Standard errors of sRSFC matrices were displayed in Supplementary Fig. 1). The absolute strength of sRSFC averaged across all connections significantly decreased in a monotonic manner as the isoflurane concentration increased (Fig. 2B, One-way ANOVA: F = 8.48, p < 10^−6^). However, the sRSFC spatial patterns remained similar across different consciousness states (Fig. 2C), reflected by a high spatial correlation coefficient between sRSFC matrices averaged across all conditions (mean (±SD) = 0.75 (±0.07)).

As largely unaltered sRSFC patterns did not seem to be able to explain dramatic behavioral change from wakefulness to unconsciousness, we further examined dynamic whole-brain connectivity patterns across all conditions. rsfMRI data at each condition were segmented using a sliding window (window length = 60 s, step size = 5 s, TR = 1 s). For each window, RSFC between each pair of ROIs was calculated using the Pearson correlation coefficient between the ROI time courses. This step yielded the RSFC matrix (i.e. a whole-brain connectivity pattern) for the window. RSFC matrices for all windows at the awake state and five anesthetic depths across all animals were then pooled together and clustered into five whole-brain connectivity patterns using k-means clustering (Manhattan distance, k = 5). These five temporarily alternating but spatially repeatable connectivity patterns were then ranked according to their spatial similarity (Manhattan distance) to the structural connectivity map generated based on axonal projection densities (Oh et al., 2014) (Fig. 3).

**Figure 3.**
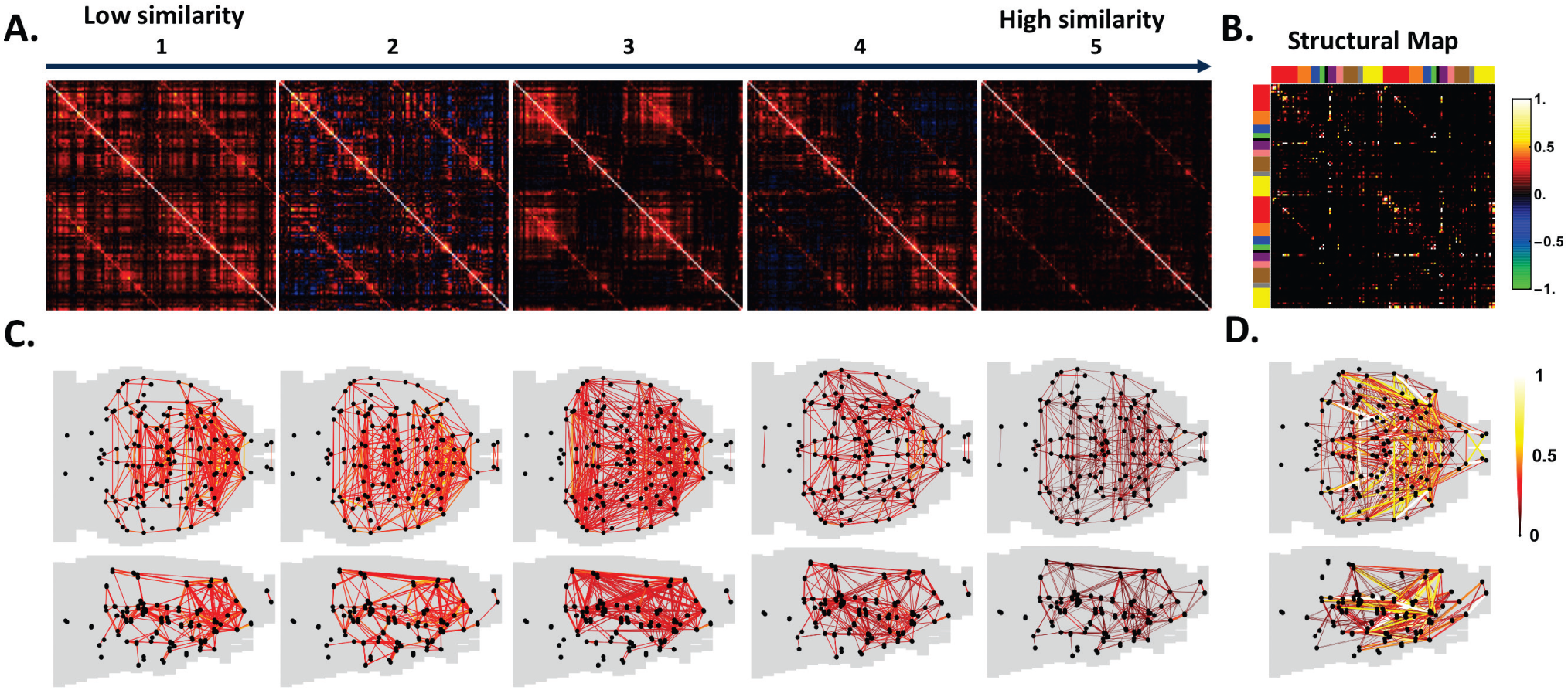
Occurrence of dRSFC patterns exhibited strong bias to different consciousness levels. Five dRSFC patterns derived from k-means clustering across all conditions. (A) 5 recurring dRSFC patterns ranked by their spatial similarity to structural connectivity (B). 134 ROIs were grouped by the hemisphere and brain system in the same way for Fig. 2. (C) Axial and sagittal views of all dRSFC patterns. (D) Axial and sagittal views of anatomical connectivity.

These dynamic connectivity patterns (referred to as dRSFC) were considerably different from the sRSFC matrices, suggesting that the brain network connectivity displayed dynamic changes that were not revealed by the conventional correlation analysis based on the stationarity assumption. In addition, all five dRSFC patterns recurred in all conditions (Standard errors of these dRSFC patterns were displayed in Supplementary Fig. 2). These results collectively indicate that whole-brain connectivity patterns contained several quasi-stable states that were spatially repeatable but temporally alternating from the awake state into anesthetized states.

We assessed the occurrence rate of all five dRSFC patterns at each conscious condition to understand the relationship of dRSFC patterns to different consciousness levels. Although all five dRSFC patterns shown in Fig. 3 were recurring in all conditions, their occurrence rate exhibited strong bias toward different consciousness levels (Two-way ANOVA: F = 17.7, p < 10^−13^ for the effect of dRSFC patterns; F = 9.46, p < 10^−25^ for the interaction between dRSFC patterns and consciousness levels). Intriguingly, we found that dRSFC patterns 1 and 5 (Fig. 4A), which were the least and most similar to the structural connectivity map, respectively, had reversed occurrence probability from the awake to unconscious states (Fig. 4C). dRSFC pattern 1 was more likely to occur when animals were at higher consciousness level and the occurrence probability of this state decreased dramatically as the anesthetic depth increased, approaching zero at the isoflurane concentration of 3.0% (One-way ANOVA: F = 10.6, p = 10^−7^). Conversely, dRSFC pattern 5 had a higher occurrence rate when animals became less conscious, and was the predominant connectivity pattern at the deepest anesthetic level (One-way ANOVA: F = 12.2, p = 10^−9^).

**Figure 4.**
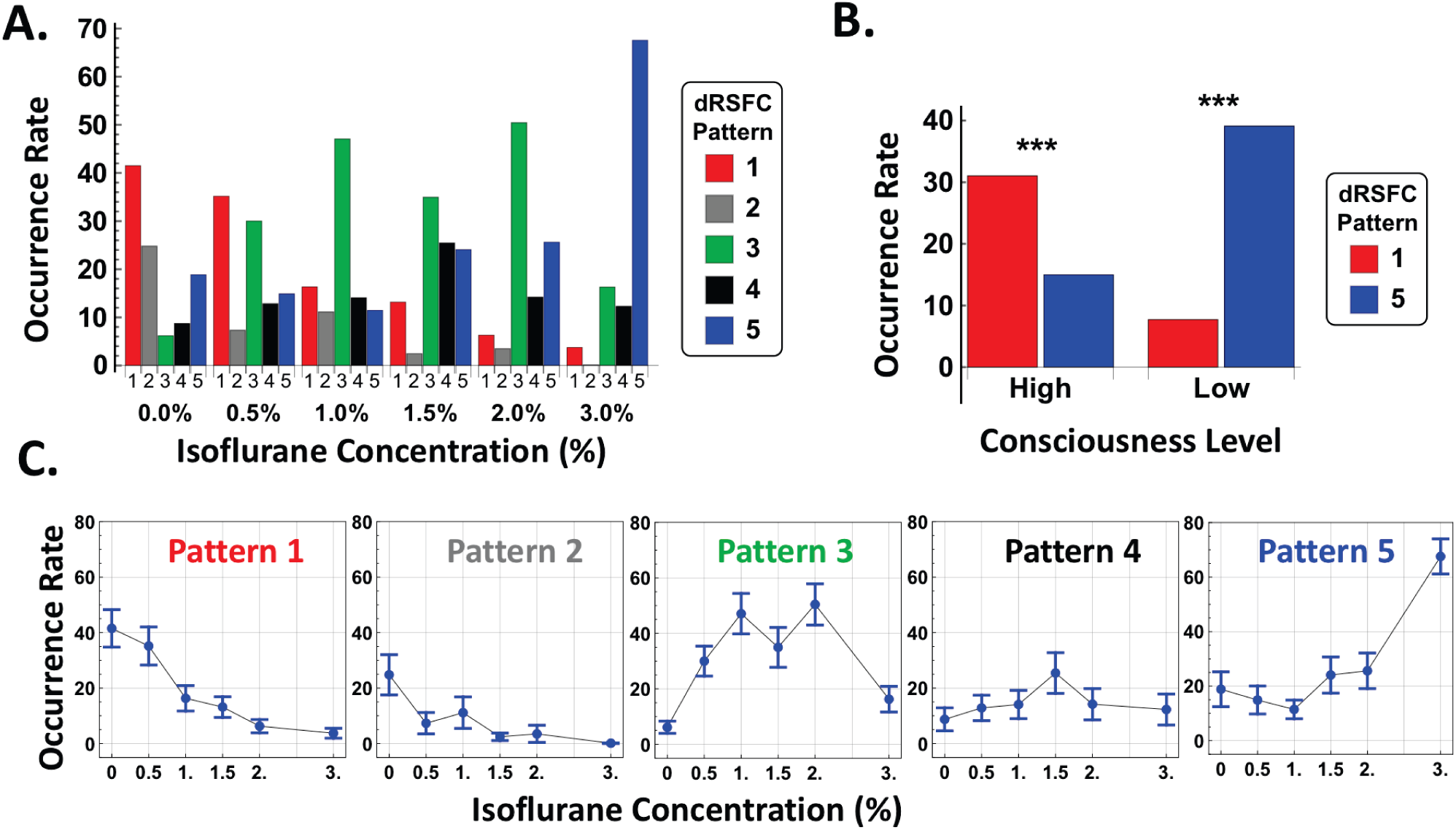
dRSFC pattern 3 represents a transition state between the conscious and unconscious states. (A) Occurrence rate for all five dRSFC patterns at each anesthetic dose. 0.0% indicates the awake state. (B) Occurrence rates of patterns 1 and 5 at two consciousness levels grouped based on the behavioral test result. Isoflurane concentrations 0.0%, 0.5% and 1.0% were assigned to the group of high consciousness level, and 1.5%, 2% and 3% were assigned to the group of low consciousness level. The occurrence probability between the two dRSFC patterns was statistically different for both consciousness levels (p<0.001 for high consciousness level and p<10^−8^ for high consciousness level). (C) Occurrence rate as a function of anesthetic depth for each dRSFC pattern, bars: S.E.M.

To further elucidate this bias in occurrence rate, all six conditions (one awake and five anesthetic conditions) were grouped into two consciousness levels based on the LORR assessment (Fig. 1B), with conditions of isoflurane 0% (i.e. wakefulness), 0.5% and 1% assigned to the high consciousness group, and isoflurane 1.5%, 2% and 3% assigned to the low consciousness group. Fig. 4B illustrates a statistically significant difference between the occurrence rate of dRSFC patterns 1 and 5 during the two consciousness levels (Two-sample t-test: p < 10^−3^ for the high consciousness level, p < 10^−8^ for the low consciousness level).

Interestingly, in addition to two brain connectivity states (dRSFC patterns 1 and 5) that were respectively characteristic to high and low consciousness levels, we observed a third connectivity state (dRSFC pattern 3) whose occurrence rate had a bell-shape distribution centered at 1.5% isoflurane (One-way ANOVA: F = 8.28, p < 10^−6^), the dose delivered when rats transitioned from the conscious state into the unconscious state (see Fig. 4C). This result suggests that this dRSFC pattern may represent a critical transitional state that occurs when the consciousness state undergoes dramatic changes at the behavioral level (Fig. 1).

In order to understand the possible transition role of dRSFC pattern 3 in switching between conscious and unconscious states, the temporal sequence of brain connectivity states for all rsfMRI windows were extracted. Each brain connectivity state was represented by the dRSFC pattern that the rsfMRI window belonged to. We calculated the state transition matrix to measure the transition probability between each two states (Fig. 5A, self-transitions were excluded). The transition probability indicates that State 1 was more likely to transition to state 3 (42% chance) than any other states, and so was State 5 (60% chance). Relative to random transitions permuted between all states while maintaining the transition probability, the preference of these transition pathways was statistically significant (State 1 to State 3: Chi-square test: χ = 41.7, p < 10^−7^; State 5 to State 3: Chi-square test: χ = 77.6, p < 10^−15^). In addition, state 3 was most likely to transition to States 1 and 5 (78% chance combined, Chi-square test: χ = 66.8, p < 10^−12^). Moreover, transitions between state 1 and state 5 were predominantly through two pathways: 1) transition through state 3, or 2) direct transition (combined 95.8% chance, Fig. 5B). Taken together, these results indicate that in addition to direct transitions between dRSFC patterns 1 and 5, dRSFC pattern 3 might represent an intermediate connectivity state between the conscious and unconscious states.

**Figure 5.**
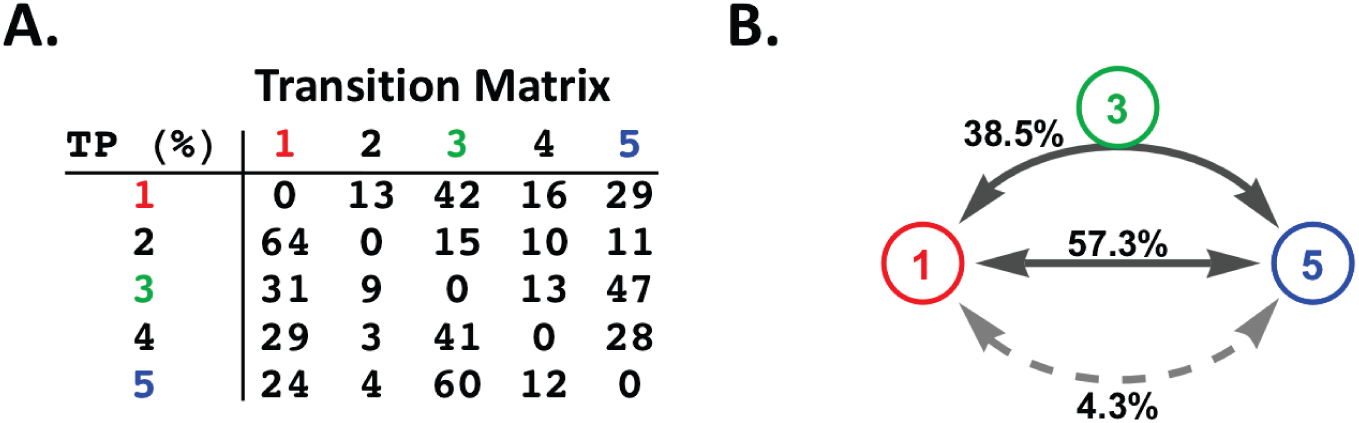
dRSFC patterns displayed distinct organizational architecture. State transitions. (A) Transition matrix between five brain connectivity states. Self-transitions were not counted. (B) Occurrence probability of different transition pathways between brain connectivity states 1 and 5. The upper line indicates that the transition path was through brain connectivity state 3. The lower line indicates the transition path was through either brain connectivity states 2 or 4, or both. The middle line suggests a direction transition between brain connectivity states 1 and 5.

To further understand the organizational architecture of dRSFC patterns of interest (patterns 1, 3 and 5), we estimated their connectivity strength distribution, the relationship between strength and connection distance, robustness, as well as the topological organization (Figs. 6 and 7). Among all three patterns, Pattern 1 displayed the most heterogeneous distribution in connectivity strength (Fig. 6A). This result is consistent with the notion that the awake state is associated with a richer functional repertoire (Barttfeld et al., 2015; Hudetz et al., 2015). The averaged absolute connectivity strength as a function of physical distance between ROIs was shown in Fig. 6B for all three dRSFC patterns. In general, all three connectivity patterns showed a trend of decreasing connectivity strength for longer-distance connections. However, State 3 displayed a ‘re-bound’ of RSFC strength for certain long-distance connections.

**Figure 6.**
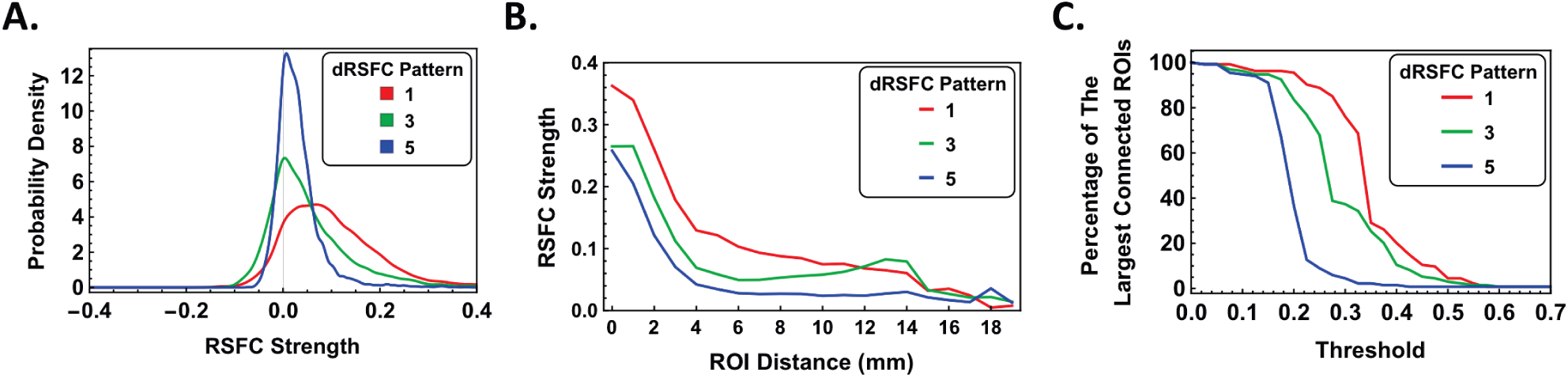
Connectivity characteristics of dRSFC networks. (A) Distribution of RSFC strength for dRSFC networks 1, 3 and 5. (B) Average absolute connectivity strength as a function of physical distance of all connections. (C) The maximal percentage of connected ROIs at different RSFC threshold for all dRSFC networks.

**Figure 7.**
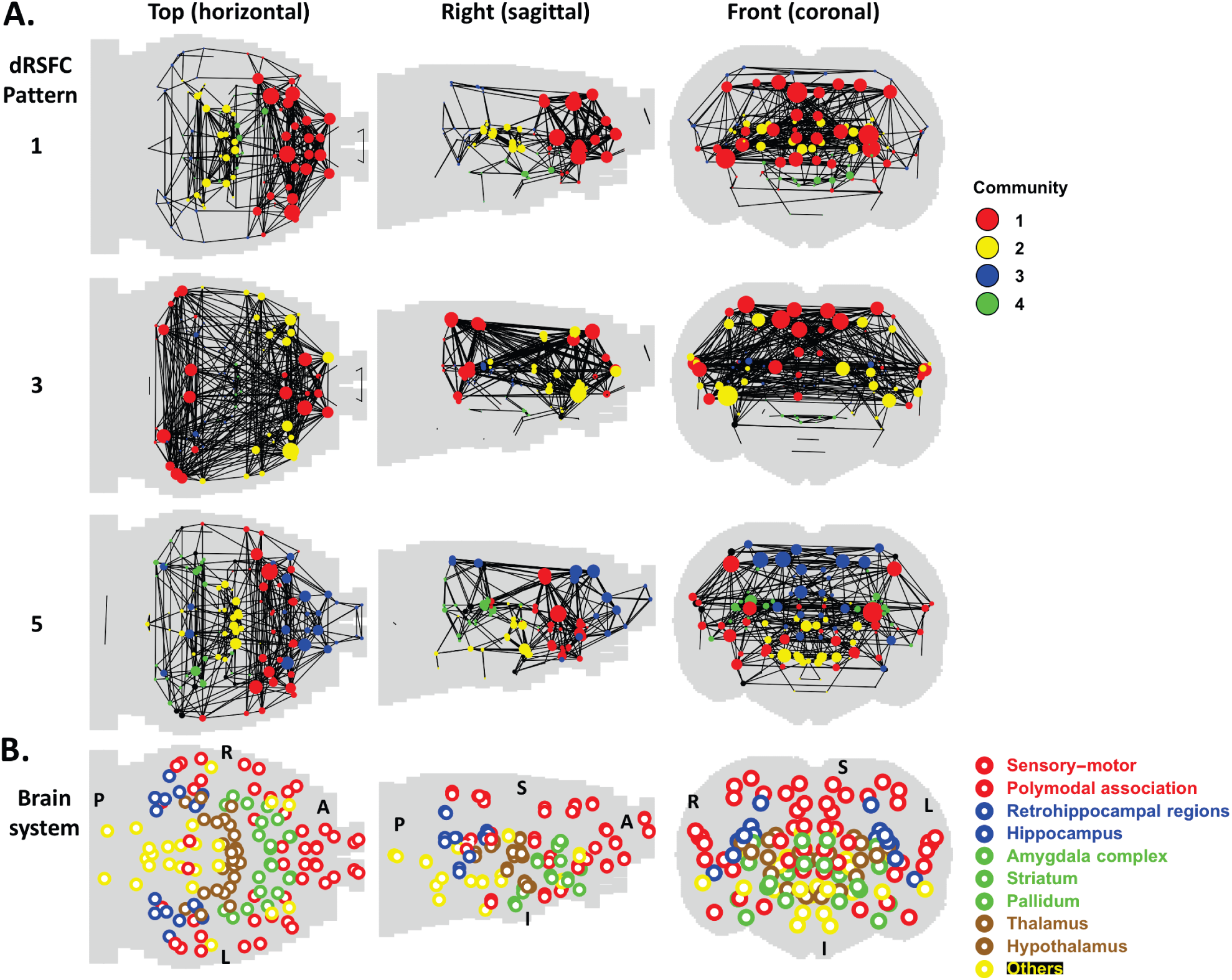
Characteristic dRSFC patterns were robust regardless of the clustering number. Community structures of dRSFC networks 1, 3 and 5 (at the graph density = 0.05).(A) Communities were color labeled. The sphere size represents the degree of the ROI. (B) The nine brain anatomical systems were displayed in four colors for the reference purpose. A, anterior; P, posterior; L, left; R, right; S, superior; I, inferior.

To assess the robustness of dRSFC networks, we calculated the percentage of ROIs of the largest connected component relative to the total ROI number at different RSFC strength threshold (Fig. 6C). In this analysis, two ROIs were deemed disconnected if the RSFC strength between them was below the threshold. This analysis can highlight the impact of losing weak connections on the network. For all three dRSFC networks, the whole-brain network began to fall into unconnected components when the threshold was elevated to a certain level. Among these three dRSFC networks, the robustness decreased in the order of network 1, 3 and 5, reflected by a gradually smaller threshold needed to fragment the network.

To examine the topological organization of dRSFC networks, we calculated the community structure of the three dRSFC networks under the same graph density (0.05). Pattern 1 primarily consisted of two communities (Fig. 7A), with one community (red) containing the sensory-motor, polymodal association and striatum systems, and the other community (yellow) containing the thalamus and hippocampus. dRSFC network 3 was characterized by three communities. The first community (red) contained brain regions in the sensory-motor, polymodal association and retrohippocampal systems. The second community (yellow) included the sensory-motor, polymodal association and striatum systems.

The third community (blue) contained part of thalamus and hippocampus. State 5 had four representative communities. Community 1 (red) included the sensory-motor, polymodal association, striatum and pallidum. Community 2 (blue) contained the sensory-motor and polymodal systems. Community 3 (yellow) included hypothalamus and a small portion of thalamus. Community 4 (green) included thalamus, hippocampus, hippocampus and retrohippocampal regions.

To examine the possible influences of the clustering number selected during k-means clustering (k=5), we also obtained results using different clustering numbers (See Supplementary Fig. 3 for the case of k = 3, and Supplementary Fig. 4 for the case of k = 7). In both cases, highly consistent dRSFC spatial patterns and occurrence preference in relation to consciousness states were revealed. At k = 3, three dRSFC patterns respectively coincided with patterns 1, 3 and 5 in the case of k=5 in terms of spatial patterns (the spatial correlations between the corresponding dRSFC patterns at k=3 and k=5 were 0.957, 0.996 and 0.917, respectively) and occurrence probability at different consciousness levels. At k = 7, the same three patterns were also identified (patterns 2, 6 and 7, the spatial correlations with dRSFC pattern 1, 3 5 at k = 5 were 0.948, 0.974 and 0.981, respectively). Taken together, these results suggest that characteristic dRSFC patterns identified were robust regardless of the clustering number.

## Discussion

The brain is an intrinsically dynamic system characterized by a variety of spatiotemporal connectivity patterns (Hutchison et al., 2013a). These functional connectivity patterns can provide important information for differentiating healthy and clinical populations (Anand et al., 2005; Carter et al., 2012; Greicius et al., 2007; Greicius et al., 2004; Hunter et al., 2012; Kennedy et al., 2006; Lowe et al., 2002; Lustig et al., 2003; Mayer et al., 2011; Tian et al., 2006; van Meer et al., 2012; Whitfield-Gabrieli et al., 2009), or can act as a ‘fingerprint’
 capable of accurately identifying individual subjects from a large group (Finn et al., 2015). An intriguing question that remains elusive is how modulations of the level of consciousness are reflected in these dynamic connectivity patterns. In the present study, we investigated the dynamics of brain network connectivity that are associated with consciousness and unconsciousness at the behavioral level. We confirm the presence of several quasi-stable whole-brain connectivity states that dynamically alternated from the awake state into anesthetized states. In particular, two brain connectivity patterns were identified to be characteristic to conscious and unconscious states, respectively. In addition, they exhibited the opposite similarity to the structure of anatomical connectivity. Interestingly, we also found a brain connectivity state that plays an important role in transitions between conscious and unconscious states. Taken together, all these findings provide compelling neuroimaging evidence that contributes to the understanding of dynamic network changes across consciousness levels. These results suggest that specific brain connectivity patterns with distinct spatial similarities to anatomical connectivity might be characteristic to different states of consciousness.

### sRSFC strength decreased, but the connectivity pattern was maintained across conscious conditions

First, we calculated the widely used sRSFC between ROIs at the awake state and five graded anesthetic depths induced by increasing concentrations of isoflurane (Fig. 2). Although the strength of sRSFC monotonically decreased with increasing isoflurane dose, we found that the spatial pattern of sRSFC generally remained similar across all conditions. This may be partially due to a global loss of information exchange under anesthesia, resulting in reduced RSFC but maintaining a similar whole-brain connectivity pattern (Tononi, 2004). This finding is also consistent with a recent study reporting resembled connectivity patterns during wakefulness, anesthesia and sleep based on electrocorticographic (ECoG) recordings from large cortical areas of macaque monkeys. (Liu et al., 2014).

### Separate dRSFC patterns were characteristic to different states of consciousness

Similar sRSFC patterns across different consciousness states can hardly explain the dramatic behavioral change from wakefulness to AIU (Keilholz, 2014). This is likely because AIU causes dynamic changes in brain function, while sRSFC data analysis can only provide averaged RSFC over the data acquisition period, losing all dynamic information. Therefore, to explore the possible link between RSFC and consciousness at the behavioral level, we used a dynamic rsfMRI approach combining the sliding-window method and k-means clustering to reveal the brain’s dRSFC patterns across all conscious conditions. Based on this analysis, we derived five brain connectivity patterns (Fig. 3) that were spatially repeatable and alternately exhibited during all states of consciousness, supporting the notion that brain network connectivity patterns contain dynamic but quasi-stable states (Allen et al., 2014; Liang et al., 2015a; Liu and Duyn, 2013; Majeed et al., 2011).

Remarkably, even though all five brain connectivity patterns recurred in all states of consciousness, separate dRSFC patterns exhibited strong bias to different consciousness levels, reflected by their distinct occurrence preference at different isoflurane doses (Fig. 4). Specifically, we identified a dRSFC pattern (pattern 5) that displayed high occurrence rate when the consciousness level is low, and became the dominant connectivity pattern at the highest isoflurane concentration. Conversely, another dRSFC pattern (pattern 1) frequently recurred when the consciousness level was high, and its occurrence rate dropped exponentially after the animal lost consciousness. Such results were observed regardless of the clustering number (k). Taken together, our results suggest that dRSFC reveals connectivity patterns that are characteristic to the conscious and unconscious states, respectively, and thus can link dynamics of brain network connectivity to consciousness at the behavioral level (Figs. 3 and 4).

### dRSFC patterns characteristic to the conscious and unconscious states contained distinct similarities to anatomical connectivity

Intriguingly, the dRSFC pattern that was strongly associated with the unconscious state was the most similar to the structural map among all five dRSFC patterns. This result is consistent with a recent finding reported by Barttfeld and colleagues (Barttfeld et al., 2015). In that study, it was suggested that functional connectivity observed during unconsciousness arises from “a semirandom circulation of spontaneous neural activity along fixed anatomical routes”, and thus that the brain’s functional connectivity pattern during unconsciousness is dictated by the structure of anatomical connectivity. Additionally, the same study showed that the conscious state contained a richer repertoire of functional configurations, although there was no investigation of a detailed relationship between these configurations and the conscious state (Barttfeld et al., 2015). In the present study, we further identified a specific connectivity pattern (dRSFC pattern 1) that was strongly associated with the conscious state. More interestingly, we found that this state was the most dissimilar from the structural map and displayed the most heterogeneous distribution in connectivity strength, indicating that information communication between different brain regions in this connectivity state is far more complicated than the restriction of anatomical connections.

The dominance of these two connectivity patterns in the conscious and unconscious states, respectively, was further confirmed by grouping all six conditions into two consciousness levels: high consciousness (wakefulness to 1.0% isoflurane) and low consciousness (1.5% to 3.0% isoflurane). We showed that the occurrence rates of these two dRSFC patterns were significantly different between the two consciousness levels (Fig. 4B). Taken together, these results suggest that specific dRSFC patterns as well as their spatial similarities to structural connectivity might be characteristic to different states of consciousness.

### Brain connectivity state during the transition from wakefulness to unconsciousness

Another interesting finding is that a brain connectivity state (dRSFC pattern 3) was dominant at the isoflurane dose regime where animals were behaviorally determined to be losing consciousness, suggesting a possible relation to the transitional phase from wakefulness to unconsciousness (Fig. 4C). To confirm this transition role, we studied the state transitions, especially between dRSFC patterns 1, 3 and 5 (Fig. 5). Firstly, transitions between these three states dominated all possible state transitions, even after considering the difference in occurrence rates. Secondly, transitions show particular preference between state 1 and 3, as well as between state 3 and 5. Thirdly, we found two typical transition pathways between state 1 and 5: 1) transition in a direct way and 2) transition through state 3. Taken together, these results indicate that in addition to direction transitions between state 1 and 5, brain connectivity state 3 likely plays a transitional role between conscious and unconscious states.

### Potential limitations

#### Isoflurane concentrations received by animals were lower than concentrations delivered

Our data indicate that loss of consciousness was induced when 1.5% isoflurane was delivered through a nosecone. This dose was higher than typical isoflurane doses inducing LORR reported in the literature (0.7%–0.8%), in which animals were either intubated or were placed in an enclosed chamber (Hudetz et al., 2011; MacIver and Bland, 2014). This apparent contradiction most likely resulted from the difference in the setup for isoflurane delivery. In our study, we could not intubate the animals as they were also imaged at the awake state in the same scanning session and intubation will unavoidably cause pain. Therefore, a nosecone was used for isoflurane delivery. As a result, the actual isoflurane concentrations in the animal were considerably lower than the concentrations delivered as air can easily entrain through the nosecone and dilute the anesthetics. However, it has to be noted that the goal of the present study is not to investigate the connectivity pattern at a specific isoflurane concentration. Instead, our goal is to use graded isoflurane concentrations to obtain different consciousness levels and examine the dynamic functional connectivity patterns in these states. Therefore, what is more important is to determine the animal’s conscious state during each rsfMRI scan. In our study, the animal’s consciousness level during each rsfMRI scan was accurately measured using the LORR test outside of the scanner in the exact manner as the imaging experiment (Fig. 1). As a result, even though the isoflurane concentrations in the animal were lower than those delivered and not precisely determined, it should not affect any conclusions of the present study.

#### Potential impact of different physiological conditions between conscious states on imaging results

It is well known that anesthesia can considerably alter physiological conditions in animals. It is likely that different physiologic conditions between the awake and anesthetized states can affect the connectivity pattern (Birn et al., 2008; Chang et al., 2009). However, our previous study suggests that influences of physiologic states on RSFC in animals are minor due to several reasons (Liang et al., 2015a). First, it has been shown that the relative contributions from cardiac and respiratory noise to the rsfMRI results were 1% and 5%, respectively, in the rat (Kalthoff et al., 2011), which are substantially smaller than the impact of physiologic noise on human rsfMRI data (Birn et al., 2008; Chang et al., 2009). This is at least partially because the breathing and heart beat rates are significantly higher in rats than humans. Since we filtered out high-frequency signals by setting the threshold of the low-pass filter at 0.1 Hz for both awake and anesthetized rats, the impact of respiratory and cardiac effects on RSFC was greatly diminished. Second, in image preprocessing we regressed out the signals from the white matter and ventricles to further minimize the influences of physiologic noise (Chang and Glover, 2009). Third, difference in physiological conditions might induce steady-state changes in functional connectivity at each anesthetic dose. However, it is unlikely to cause systematic dynamic RSFC patterns found in the present study, particularly considering that all dRSFC patterns recurred in all anesthetic conditions. In fact, in our previous study (Liang et al., 2015a) we calculated the correlation between the BOLD signals from the white matter and ventricles and the time series of each individual voxels. The spatial pattern of this correlation departed far away from any dynamic connectivity patterns (awake or anesthetized) reported in that study, suggesting minimal influences from physiologic noise on dynamic RSFC in rats. These results collectively indicate that difference in physiological conditions cannot account for any major effects of dRSFC patterns in the present study.

#### Impact of motion

Another potential pitfall is that dRSFC patterns could be attributed to the difference in the motion level at different anesthetic depths. Motion decreased as the anesthetic level increased, as shown in Supplementary Fig. 5a. In addition, the motion level of dRSFC pattern 1 was higher than those of dRSFC patterns 3 and 5 (Fig. 1b), presumably because dRSFC pattern 1 was strongly biased by the awake state, while patterns 3 and 5 were more dominant in unconscious states. To rule out the possibility that characteristic dRSFC patterns identified in the present study were motion artifacts, we selected a subgroup of animals (n=15), whose movement level was minimal in all rsfMRI scans. Three dRSFC patterns with the same occurrence probability in relation to the consciousness level were also identified in this subset of data (Supplementary Fig. 6, Patterns 3, 4 and 5 respectively correspond to Patterns 1, 3 and 5 in the full dataset. Patterns 1 and 2 had extremely low occurrence rate, suggesting that they are most likely outliers). Notably, the motion levels in dRSFC patterns 3, 4 and 5 in this low-motion dataset were comparable (Supplementary Fig 6c), indicating that characteristic dRSFC patterns revealed in the present study were not attributed to movement.

Other limitations include that the structural connectivity was derived from the mouse brain atlas. Although highly similar, there is still subtle difference in the shape between the rat and mouse brains. It also has to be noted that the findings of the present study can only be applied to isoflurane. To generalize these findings, it is necessary to conduct a similar study using different anesthetic agents.

## Acknowledgement

We would like to thank Dr. Pablo Perez for his technical support, and Ms Lilith Antinori for editing the manuscript. Dr. Nanyin Zhang is supported by R01MH098003 from the National Institute of Mental Health and R01NS085200 from the National Institute of Neurological Disorders and Stroke.

